# Intercostal Space Prediction Using Deep Learning In Fully Endoscopic Mitral Valve Surgery

**DOI:** 10.1101/710996

**Authors:** Rafik Margaryan, Daniele Della Latta, Giacomo Bianchi, Nicola Martini, Gianmarco Santini, Dante Chiappino, Marco Solinas

**Affiliations:** Via Aurelia Sud 303, Massa, Tuscany, 54100, Italy

## Abstract

**Objective:** About 10 million people in Europe suffer from mitral valve incompetence. Majority of these entity is mitral valve prolapse in developed countries. Endoscopic mitral valve surgery is a relatively new procedure and preparation in the right intercostal space are crucial for success completion of the procedure. We aimed to explore clinical variables and chest X-rays in order to build most performant model that can predict the right intercostal space for thoracotomy.

**Methods:** Overall 234 patients underwent fully endoscopic mitral valve surgery. All patients had preoperative two projection radiography. Intercostal space for right thoracotomy was decided by expert cardiac surgeons taking in consideration the height, weight, chest radiography, anatomical position of skin incision, nipple position and the sex. In order to predict the right intercostal space we have used clinical data and we have collected all radiographies and feed it to deep neural network algorithm. We have spitted the whole data-set into two subsets: training and testing data-sets. We have used clinical data and build an algorithm (Random Forest) in order to have reference model.

**Results:** The best-performing classifier was GoogLeNet neural network (now on we will reffera as Deep Learning) and had an AUC of 0.956. Algorithm based on clinical data (Random Forest) had AUC of 0.529 using only chest x-rays. The deep leaning algorithm predicted correctly in all cases the correct intercostal space on the training datasest except two ladies (96.08% ; with sensitivity of 97.06% and specificity 94.12 %, where the Random Forest was capable to predict right intercostal space in 60.78% cases with sensitivity of 93.33% and specificity 14.29 % (only clinical data).

**Conclusion:** Artificial intelligence can be helpful to program the minimally invasive cardiac operation, for right intercostal space selection for thoracotomy, especially in non optimal thoraxes (example, obese short ladies). It learned from the standard imaging (thorax x-ray) which is easy, do routinely to every patient.

## Introduction

Mitral valve regurgitation (MR) is the second most frequent indication for valve surgery in Europe and it continues to grow. The prevalence of mitral valve prolapse is oscillating between 1 to 2.5 % in developed countries[1]. Usually these are around 60 years old patients who expect to have good quality of life and life expectancy is over 15 - 20 years. Interest in mitral valve repair in fully endoscopic settings is growing [2]. Many groups have reported their results using different types of endoscopic setup [3,4]. However due to large variation of chest dimensions and anatomy there is a discrepancy in choosing the correct thoracotomy intercostal space for given patient. Our experience in mitral valve surgery[5] showed feasibility in the forth intercostal space when using rib spreading and central cannulation (direct vision guided surgery). Many authors choose third intercostal space and some others [4] fourth intercostal space as a default for endoscopic mitral valve surgery (video guided surgery). Intercostal space correct prediction is important of terms of feasibility, short cardio-pulmonary bypass time, short aortic cross clamping (one of most important variables) and overall intervention time. Recently, many approaches for medical imaging analysis have applied deep convulutional networks or ‘deep learning’ instead of engineering image features[6]. Deep learning is an example of representation learning (without pre-specification of discreiminative features, it learnes for raw data). There are some unwritten rules in different institutions which give some intraoperative tips on how to determine the right intercostal space for thoracotomy; for example, the middle of the thorax measured by both hands (in our institution it is called Farneti’s rule, see Figure 1. It is difficult to standardize the approach for given thorax, and it is operator dependent and not reproducible, therefore we attempt here to find a way to predict intercostal space in given thorax.

**Fig 1.**
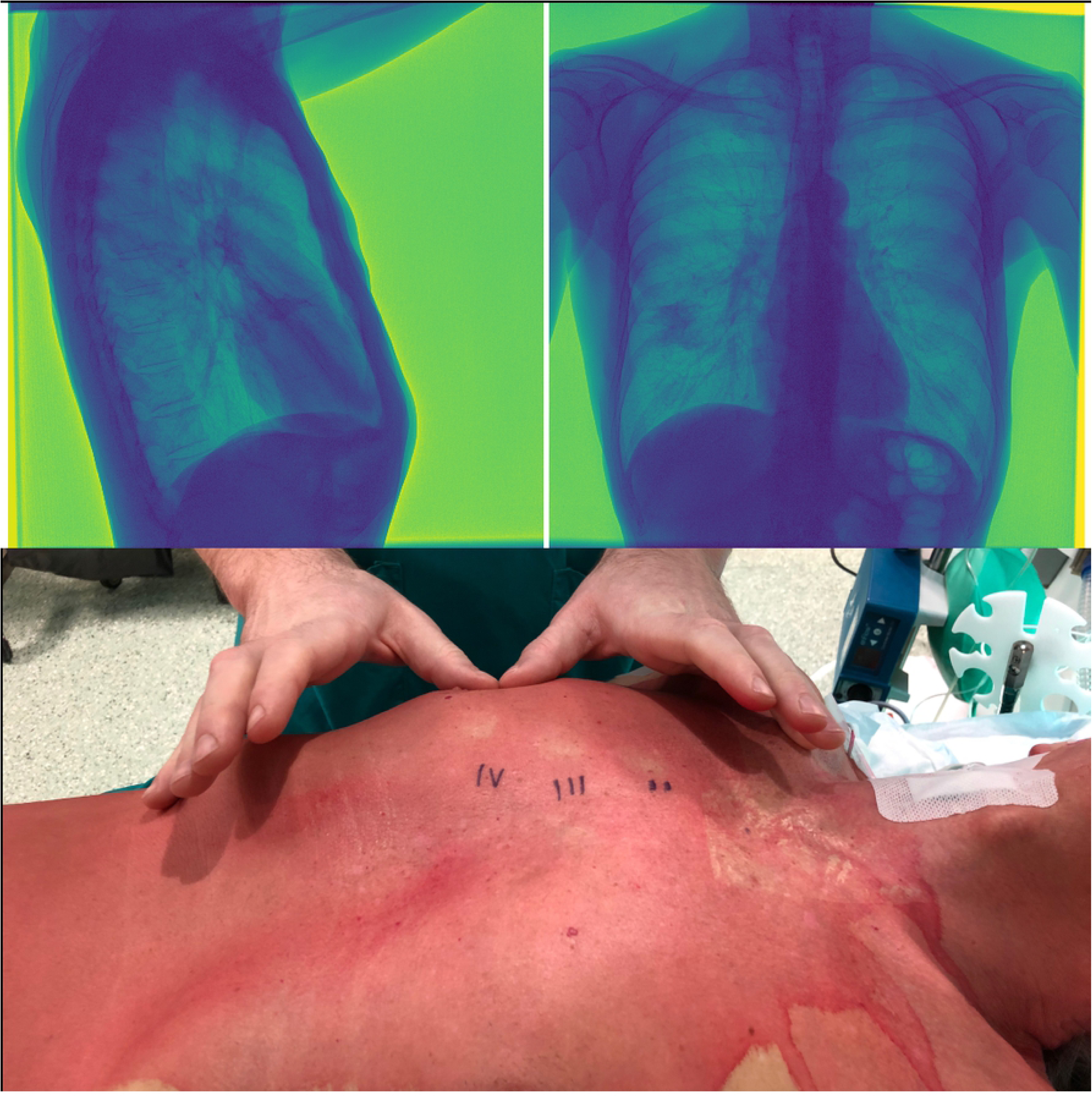
Thorax measurements. A: Anterposterior and lateral radiophraphy. B: Farnetis rule for the intercostal space determination

### Objective

We set a goal to use:

1. all relevant clinical data using one of most performant machine learning algorithms (Random Forest as one of most performant algorithms[7])
2. Deep learning in order to predict intercostal space using only chest radiography itself (neural networks, GoogLeNet).

## Material and Methods

### Patients’ Characteristics

234 consecutively operated patients from 2015 to 2018 underwent totally endoscopic mitral valve surgery via limited 3-4 cm right thoracotomy. All patients had preoperative chest x-ray (antero-posterior and lateral). For demographic data see Table 2. Intercostal space approach was decided upon radiography analysis by experienced cardiac surgeon. All intervention where performed by senior cardiac surgeon. All clinical data was collected prospectively. All mitral patients were forwarded to endoscopic surgery and had different levels of complexity: 1) Simple if there were two procedures (ex. annuloplasty and one/two pair of neochordae positioning) or any other combination of two surgical repair gestures (resection, sliding, commisural closure, Alfieri stitch, etc). 2) Complex procedures where we preformed three or more surgical gestures during single mitral valve repair operation. 3) All mitral valve replacement cases were considered as a complex mitral valve procedure in endoscopic settings. For complexity see Table 4.

**Table 1.**
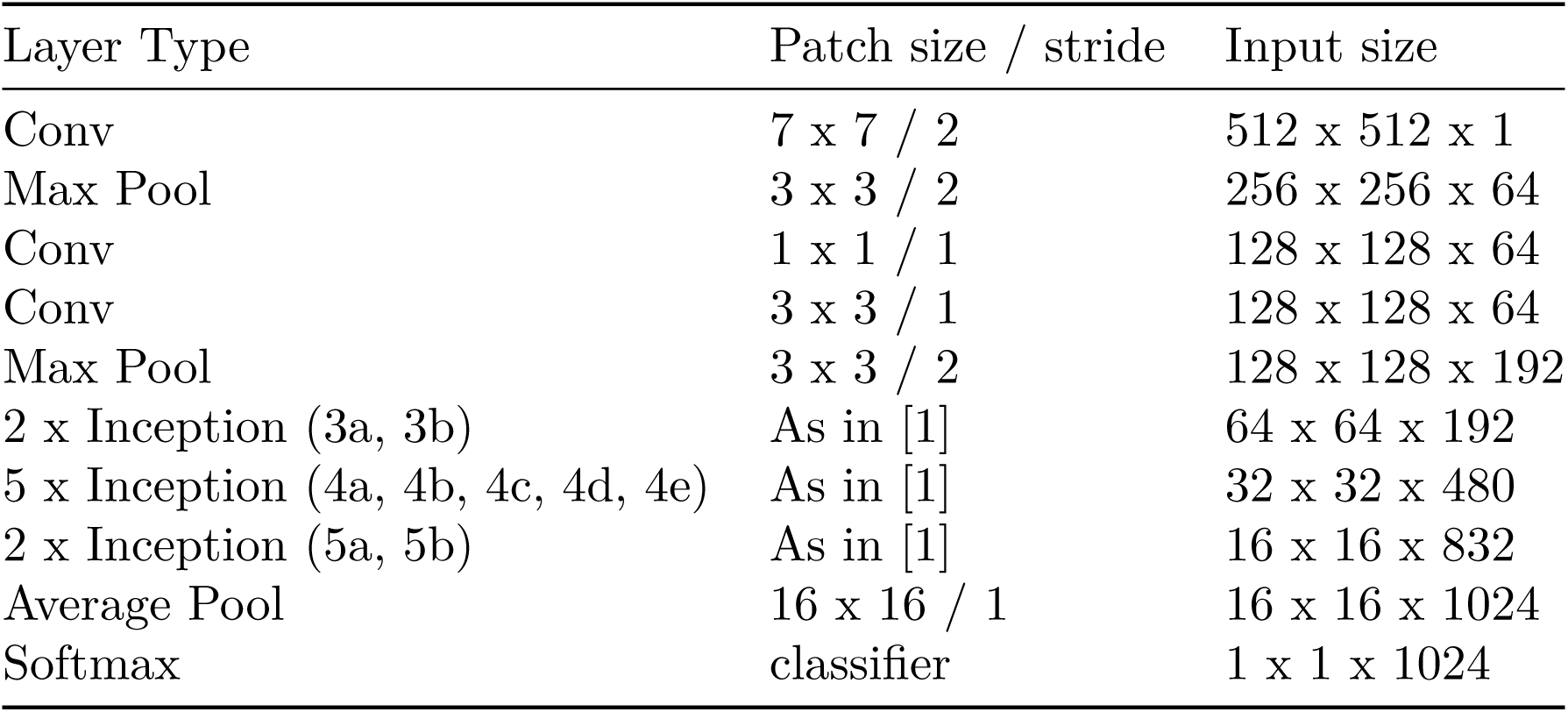
Schematic representation of the GoogLeNet network used to classify the input images. For a detailed description of the network architecture see [8].

**Table 2.** Demographics

|  | 3rd<br>N=149 | 4th<br>N=85 | p.overall |
| --- | --- | --- | --- |
| Sex: |  |  | <0.001 |
| Female | 32 (21.5%) | 42 (49.4%) |  |
| Male | 117 (78.5%) | 43 (50.6%) |  |
| Mitral_Procedure_Complexity: |  |  | 0.185 |
| No | 118 (79.2%) | 60 (70.6%) |  |
| Yes | 31 (20.8%) | 25 (29.4%) |  |
| CPB | 144 (28.2) | 143 (29.9) | 0.735 |
| Aortic_Cross_Clamping | 92.3 (20.8) | 90.6 (22.7) | 0.563 |
| Intervention_Time | 243 (48.1) | 224 (160) | 0.281 |
| Peso | 111 (429) | 133 (569) | 0.759 |
| Altezza | 170 (19.9) | 170 (9.95) | 0.936 |
| Approach: |  |  | <0.001 |
| Areolare | 111 (74.5%) | 39 (45.9%) |  |
| Mini | 38 (25.5%) | 46 (54.1%) |  |
| CorKnot: |  |  | 0.002 |
| No | 12 (8.05%) | 20 (23.5%) |  |
| Yes | 137 (91.9%) | 65 (76.5%) |  |
| MVP: |  |  | 0.002 |
| none | 0 (0.00%) | 1 (1.18%) |  |
| repair | 141 (94.6%) | 69 (81.2%) |  |
| replacement | 8 (5.37%) | 15 (17.6%) |  |
| ConversioneM: |  |  | 0.020 |
| No | 146 (98.0%) | 77 (90.6%) |  |
| Yes | 3 (2.01%) | 8 (9.41%) |  |
| ThreeD: |  |  | 0.842 |
| 2D | 136 (91.3%) | 79 (92.9%) |  |
| 3D | 13 (8.72%) | 6 (7.06%) |  |

**Table 3.** Test Table

|  | Female |  |  | Male |  |  |
| --- | --- | --- | --- | --- | --- | --- |
|  | 3rd<br>N=10 | 4th<br>N=10 | p.overall | 3rd<br>N=24 | 4th<br>N=7 | p.overall |
| RF_Prediction: |  |  | 1.000 |  |  | 1.000 |
| 3rd | 9 (90.0%) | 9 (90.0%) |  | 21 (87.5%) | 7 (100%) |  |
| 4th | 1 (10.0%) | 1 (10.0%) |  | 3 (12.5%) | 0 (0.00%) |  |
| DL_Prediction: |  |  | 0.002 |  |  | <0.001 |
| 3rd | 9 (90.0%) | 1 (10.0%) |  | 24 (100%) | 0 (0.00%) |  |
| 4th | 1 (10.0%) | 9 (90.0%) |  | 0 (0.00%) | 7 (100%) |  |

**Table 4.** Procedure by intercosatal space

|  | 3rd<br>N=149 | 4th<br>N=85 | p.overall |
| --- | --- | --- | --- |
| Annuloplasty: |  |  | 0.002 |
| No | 8 (5.37%) | 16 (18.8%) |  |
| Yes | 141 (94.6%) | 69 (81.2%) |  |
| Neochorde: |  |  | 0.002 |
| No | 49 (32.9%) | 46 (54.1%) |  |
| Yes | 100 (67.1%) | 39 (45.9%) |  |
| Alfieri: |  |  | 1.000 |
| No | 148 (99.3%) | 84 (98.8%) |  |
| Yes | 1 (0.67%) | 1 (1.18%) |  |
| Sliding: |  |  | 0.194 |
| No | 147 (98.7%) | 81 (95.3%) |  |
| Yes | 2 (1.34%) | 4 (4.71%) |  |
| Gender: |  |  | <0.001 |
| Female | 32 (21.5%) | 42 (49.4%) |  |
| Male | 117 (78.5%) | 43 (50.6%) |  |
| Chiusura: |  |  | 1.000 |
| No | 128 (85.9%) | 73 (85.9%) |  |
| Yes | 21 (14.1%) | 12 (14.1%) |  |
| Resezione: |  |  | 0.798 |
| No | 132 (88.6%) | 77 (90.6%) |  |
| Yes | 17 (11.4%) | 8 (9.41%) |  |
| Complex_MI_Procedure: |  |  | 0.185 |
| Simple | 118 (79.2%) | 60 (70.6%) |  |
| Complex | 31 (20.8%) | 25 (29.4%) |  |
| MVP: |  |  | 0.002 |
| none | 0 (0.00%) | 1 (1.18%) |  |
| repair | 141 (94.6%) | 69 (81.2%) |  |
| replacement | 8 (5.37%) | 15 (17.6%) |  |

### Endoscopic Mitral Valve Procedure

Under general anesthesia, patients were intubated using a single lumen endotracheal cannula for pulmonary ventilation. Disposable paddles for external cardiac defibrillation were placed in the right scapula and anterolateral region of the left hemi-thorax. A two- or three-dimensional transducer for intraoperative trans esophageal electrocardiography (TEE) was placed. Before the positioning the patient, the right hemi-thorax was carefully examined, and significant pre-existing asymmetries or chest deformities were recognized: that might predict complicated access via periareolar access. After patients have been positioned whit the right hemi-thorax about 15-20 degrees, marking the reference intercostal spaces and lines were cared out in order to facilitate the anatomical references when sterile field is completed (we do use sterile elastic field membrane). A right angle skin incision was performed inside the edge of the nipple-areolar complex, on its lower half circumference (hours position of 4 to 10, clockwise). In female patients we prefer to do skin incision in the inframmamary line as wide as periareolar incision (3-4 cm). Subcutaneous tissues was then sectioned with dietary through the entire length of the incision, reaching the major pectoral musculus the underlying intercostal space. Later hemostasis was performed accurately. Soft tissue retractor was used providing 360 degree atraumatic circumferential retraction and allowing maximum exposure with minimum incision size. No rib retractore was used in this approach. A 10.5 (some cases 12.5) mm trocar was then inserted in the same intercostal space as the incision on the mid-axillary line ; and two intercostal spaced lower (5th or 6th intercostal spaces) in the anterior axillary line. Femoral vessels were isolated. Systematic heparinization was achieved (HMS or simple ACT guided). At that point the femoral vessels were cannulated using Seldinger technique under direct vision. Venous cannula (Biomedicus 25 F) were placed in the right atrium using trans-esophageal echocardiography. In the case of suspected insufficient single cannula venous drainage a second cannula was placed into the jugular vain (16 F). Seldinger technique was used also for femoral arterial cannulation with Biomedicus 15-21 F cannulas depending on the weight. Cardio-pulmonary bypass was established and the patients were cooled to 34°C. After pump is started the meccanical ventilation was suspended. A pledgeted 4/0 polipropilen purse-string suture for cardioplegia and aortic venting was placed on the ascending aorta. Through the 2nd intercostal space in the anterior axillary line the Chitwood transthoracic clamp was introduced (Fehling Instruments, Karlstein, Germany). Under the video guidance the ascending aorta was clamped and cold crystalloid cardioplegic solution Custodiol HTK (Kohler Chemie GmbH, Bensheim, Germany) was administrated. The surgical instruments used were not specific for endoscopic mitral valve surgery; they are the routine minimally invasive surgery instruments. Under the videoscopy the pericardium was opened 2-3 cm above the phrenic nerve. The pericardium was retracted with three polypropylene 3/0 and three polyester 2/0 suture (above and under, respectively). The inferior retraction suture were extemporized using two ports and entrance for Chitwood clamp. With the aim of air embolism prevention CO_2_ insufflation was maintained during central part of the procedure (before atrial cavity opening and stopped after its closure and decamping).

### Mitral valve repaire

The left atrium was opened anteriorly to the right pulmonary veins and retracted using a trans-thoracic retractor (Estech, San Ramon, CA, USA or USB retractor, Belgium) positioned through the same intercostal space as the thoracotomy medially or laterally to right internal mammary artery. The mitral valve was inspected and then repaired or replaced. In the mitral valve repair the following techniques were used: 1. annuloplasty 2. leaflet resection 3. neochordal implantation 4. comisural or any other leaflet closure 5. Alfiery technique The valve competence was tested with cold saline solution after the repair was complete. In case of the mitral valve replacement mechanical or tissue valves were used. Additionally, when necessary tricuspid valve annuloplasty was carried out. In some patients atrial fibrillation the box lesion was created using Cobra System before the left atrial opening. After the procedure the atriotomy was closed with 3/0 polypropylene running suture and a ventricle vent was positioned through the atriotomy. Aim of the study was to develop an algorithm that could predict the intercostal space with highest accuracy possible. We have developed the machine learning algorithm using clinical data and some simple radiography measurements; and a deep learning algorithm that could only use the chest radiography for intercostal space prediction.

### Statstical analysis

Clinical data was collected in prospective fashion. Continuous variables where presented as mean and standard deviation and compared using non parametric and parametric test (Wilcos test, Student’s test, respective). Categorical variables where presented as percentage compared with Fisher’s exact test or *χ*^2^ test. Algorithm bases on clinical data was created using random forest tree classification algorithm. All the analysis were performed using R, Python programming languages using reproducible research rules.

### Data Acquisition and Processing

234 chest digital X-ray examinations (DX) were performed on patients enrolled for valvular surgery. X-ray images of lateral latero (LL) projections were used to train a deep convolutional neural network (DNN). All DX were taken with standing patient placed 180 cm from the X-ray tube using a digital flat detector panel with a spatial resolution of 0.139×0.139 mm^2^. The size of the image matrix is variable (2500×2500-4000×4000 px) depending on the size of the patient’s chest as well as the X-ray exposure. The potential difference (kV) and exposure current (mAs) parameters are set automatically of the X-ray scanner in order to obtain an adequate contrast independently of the variation of the attenuation of the body of each patient. The dataset was splitted into 183 images to train the DNN and 22 to test the learning. The size of all the images was readjusted to 2000×2000 with a centered crop operation so as to maintain the DNN input layer with fixed size and partially mask the background that does not provide an useful information for the classification task. Finally, to minimize the computational load for network training, the images were scaled to 512× ×512 px. Whole data was divided randomly into two parts: train dataset and test dataset. Train dataset was uesed to create the model and validate it. Test dataset was used for only external validation and never has been used for tuning or improving the algorithm.

### Network Architecture

A GoogLeNet network [8] was used to perform the classification task. The outline of the proposed architecture was reported in Table 1. The network comprises 22 trainable layers and is made by nine Inception module stacked upon each other. A single Inception module represents a combination of 1×1, 3×3 and 5×5 convolutional kernels, used in parallel with the aim to progressively cover a bigger area and extract details (1×1) as more general features (5×5). The results are then concatenated with the outcome of a max-pooling layer and passed to the next layer. The presence of 1×1 convolutions before the 3×3, 5×5 convolutions and after the pooling layer help to reduce the number of parameters in the Inception module and add more non-linearities to the network. To apply the GoogLeNet network architecture to the surgical task some implementation changes had to be made: 1) the input layer dimensions, set at 512 × 512, with a number of channels equals to one instead of three, being the inputs grayscale images; 2) in the output layer only two units were used, as we had to classify the images according the two possible locations of surgical incision; The network was designed and trained with Tensorflow [9], running for 500 epochs on a single NVIDIA Titan Xp. Stochastic gradient descent and Adam optimizer were used to minimize a categorical cross-entropy for the network parameters update.

Schematic representation of the GoogLeNet network used to classify the input images. For a detailed description of the network architecture see [8].

The size of all the images was readjusted to 2000×2000 with a centered crop operation so as to maintain the DNN input layer with fixed size and partially mask the background that does not provide an useful information for the classification task. Finally, to minimize the computational load for network training, the images were scaled to 512×512 px. To evaluate the progress of learning the accuracy was calculated between the predicted and the real “point of incision”.

## Results

### Patients’ clinical data

Cohort consisted of 234 patients, collected at the Ospedale Del Cuore of Massa, Italy. All patients underwent successful mitral valve procedure. Unfortunately there were one hospital and/or 30 days mortality event 1 (0.427%) due to post procedural acute heart failure. Mean age was 61.3 ± 11.8 and 74 (31.6%) were female patients. All patients are of Caucasian ethnicity. Forth intercostal space was more often preferred for the female patients due to the breast tissue, and third intercostal space was frequently approached in male patients (periareolar cut more often is corresponding to 3rd intercostal space, biased by preconception). Cardio-pulmonary bypass and aortic cross clamping time was not different in both groups (see Table 2). Deep learning prediction algorithm predicted the correct intercostal space for all the patients except two female patients. Random forest was able to predict only in 31 cases. It misclassified female patients and male patients, whereas the deep leaning had misclassified only female patients (see Table 3).

### Classification

Using only clinical data it was possible to achieve 60.78% accuracy (range 46.11 - 74.16 %, see Figure 3 and Table **??**). The most important factor for intercostal space prediction was weight, followed by height (see Figure 3). There were general agreement on the classification (See Table 3) There were 97.059 % agreement on 3rd and 94.118 % agreement on 4th intercostal space for deep learning algorithm and 88.235 % agreement on 3rd and 5.882 % agreement on 4th intercostal space for random forest algorithm when ignorig the gender of the patients. For specific disagreemend between taking in consideration also gender see Table 3. The best-performing classifier was GoogLeNet neural network (Deep Learning) and had an AUC of 0.9608 (see Figure 4. The deep leaning algorithm predicted correctly in all cases the correct intercostal space on the training datasest except two ladies (96.08% ; with sensitivity of 97.06% and specificity 94.12 % (see Table **??**)

**Fig 2.**
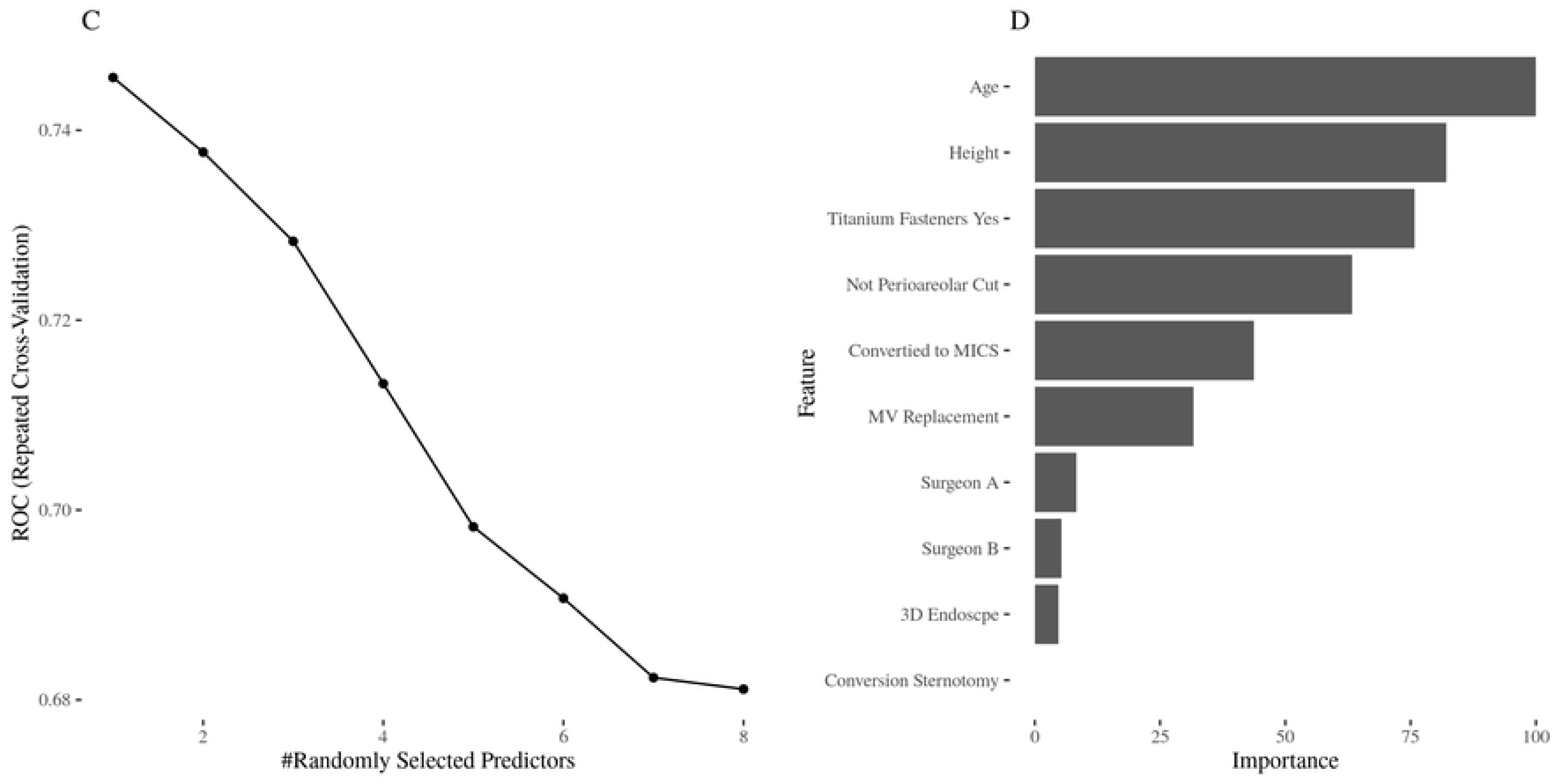
Random Forest accuracy plot. A: Radom Forest traing curve with random variables. B: Random Forest with variable importance

**Fig 3.**
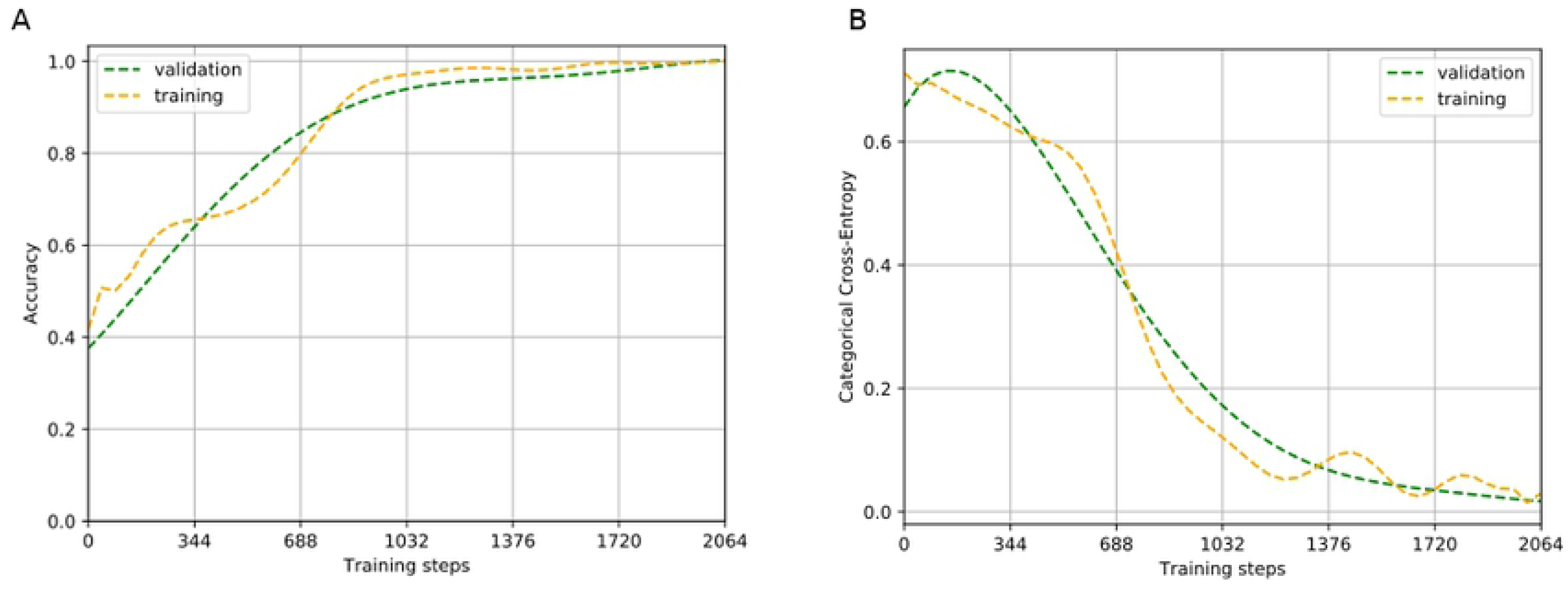
Deep Learning accuracy plot. A: Deep Learning Accuracy gain durein training. B:Deep Learning training steps C: Radom Forest traing curve with random variables. D:Random Forest with variable importance

**Fig 4.**
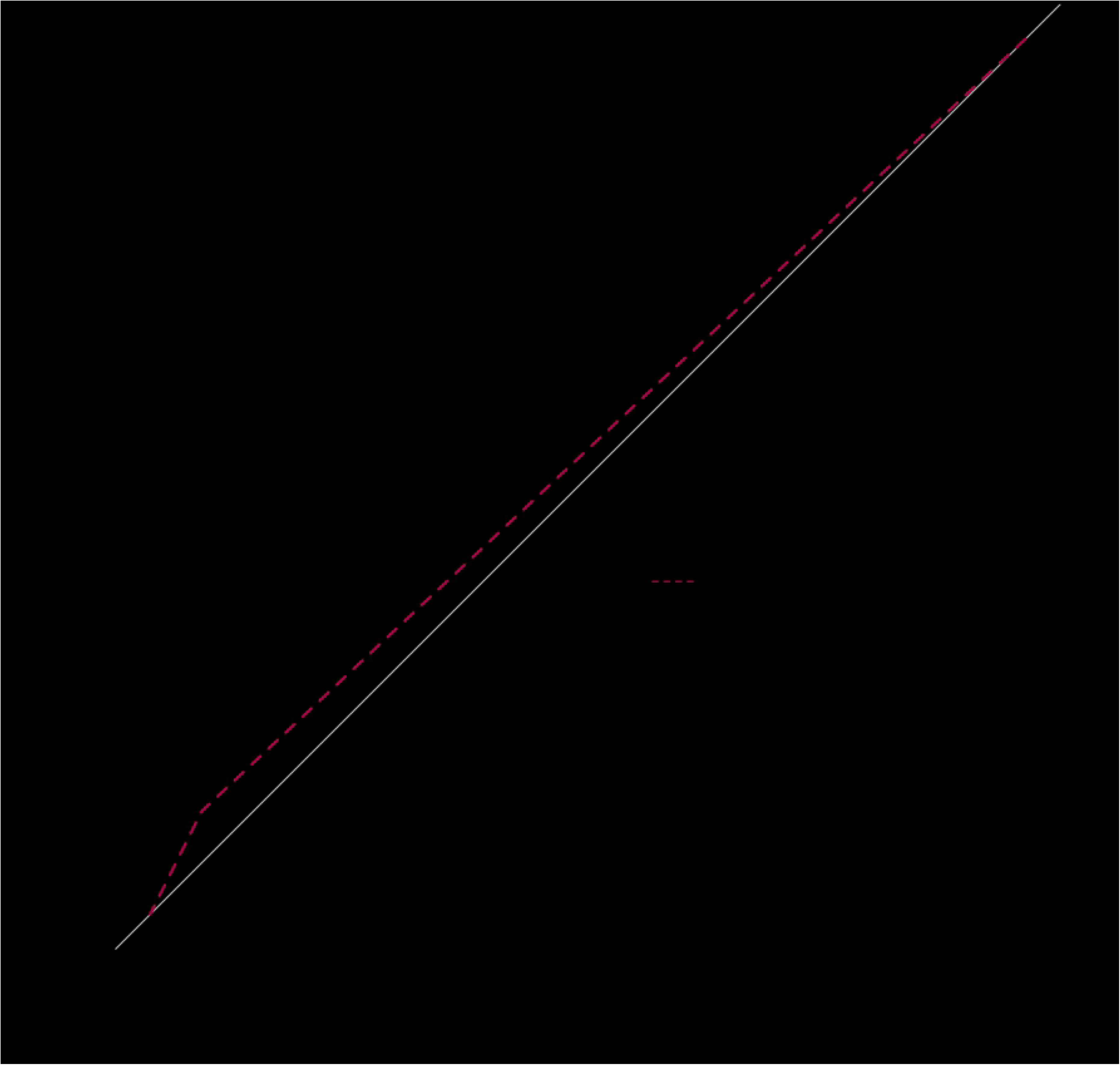
AUC curve for both algorithms

## Discussion

Mitral valve reconstructive surgery is challenging. It is well known that mitral valve repair can last more then 15 and more years [10,11]. Minimally invasive approach can shorten the hospital stay, less blood product transfusion[5]. In fully endoscopic mitral valve surgery choosing right approach in terms of intercostal space and right angle intervention itself can be performed with minimal fatigue and short CPB time and aortic cross clamping time. Wrong selection of the intercostal space can result of approach failure, therefore, converted to conventional thoracotomy or complete sternotomy. 4th intercostal space is a standard for direct vision mitral and/or other type of valve surgery [5,12].

There is a lot difficulties related to subcutaneous tissue especially when it concerns to obese patients. These are not appreciated when using only weight of the patients or other variable related to it. Big abdomen with severe amount of fat deposits shift the diaphragm in this cases towards lung apex more then it is expected from radiography when anesthetist.

Artificial intelligence was started to be applied to medicine and other scientific disciplines[13]. Very recently only the computer become fast enough in order to be fore simple application. Chest radiography is routine procedure for many institutions (if not all for them) and it is easy to do. Using simple simple two projection radiographies with sophisticated algorithm we have reached a very high accuracy. This makes us thing that there are some anatomical substrates hidden in the chest radiography which are possible to extract/make use of it using artificial intelligence.

Mentoring in minimally invasive cardiac surgery is a debated issue and it merits a very important attention [14]. Artificial intelligence can be a guide for young surgeons or new programs. We believe that this application of the artificial intelligence could help to find right approach almost for every patient, thus, every newcomer in single mitral valve disease can be addressed to endoscopic program.

## Conclusions

Artificial intelligence can be helpful to program the minimally invasive, fully endoscopic mitral operation, in order to find the right intercostal space, especially in non optimal thoraxes. It learns from the standard imaging (thorax radiography) which is easy to do routinely and it is cost effective. It is important for mentoring process.

## Author Contribution

### Conceptualization

Rafik Margaryan, Giacomo Bianchi, Daniele Della Latta

### Clinical Data

Rafik Margaryan, Giacomo Bianchi

### Radiological Data

Daniele Della Latta

### Formal Analysis

Rafik Margaryan, Daniele Della Latta

### Writing - Original Draft

Rafik Margaryan

### Writing - revew & editing

Rafik Margaryan, Giacomo Bianchi, Daniele Della Latta.

## References

1. Freed LA, Levy D, Levine RA, Larson MG, Evans JC, Fuller DL, et al. Prevalence and clinical outcome of mitral-valve prolapse. The New England Journal of Medicine. 1999;341: 1–7. doi:10.1056/NEJM199907013410101

2. Schmitto JD, Mokashi SA, Cohn LH. Minimally-invasive valve surgery. Journal of the American College of Cardiology. 2010;56: 455–462. doi:10.1016/j.jacc.2010.03.053

3. Lamelas J, Nguyen TC. Minimally Invasive Valve Surgery: When Less Is More. Seminars in Thoracic and Cardiovascular Surgery. 2015;27: 49–56. doi:10.1053/j.semtcvs.2015.02.011

4. Poffo R, Montanhesi PK, Toschi AP, Pope RB, Mokross CA. Periareolar Access for Minimally Invasive Cardiac Surgery: The Brazilian Technique. Innovations (Philadelphia, Pa). 2018;13: 65–69. doi:10.1097/IMI.0000000000000454

5. Glauber M, Miceli A, Canarutto D, Lio A, Murzi M, Gilmanov D, et al. Early and long-term outcomes of minimally invasive mitral valve surgery through right minithoracotomy: A 10-year experience in 1604 patients. Journal of Cardiothoracic Surgery. 2015;10: 181. doi:10.1186/s13019-015-0390-y

6. LeCun Y, Bengio Y, Hinton G. Deep learning. Nature. 2015;521: 436–444. doi:10.1038/nature14539

7. Breiman L. Random forests. Mach Learn. Hingham, MA, USA: Kluwer Academic Publishers; 2001;45: 5–32. doi:10.1023/A:1010933404324

8. Szegedy C, Liu W, Jia Y, Sermanet P, Reed S, Anguelov D, et al. Going Deeper with Convolutions. arXiv:14094842 [cs]. 2014; Available: http://arxiv.org/abs/1409.4842

9. Workshop on Wireless Traffic Measurements and Modeling, USENIX Association, ACM SIGMOBILE, ACM Special Interest Group in Operating Systems, ACM Digital Library. Papers presented at the Workshop on Wireless Traffic Measurements and Modeling: June 5, 2016,Seattle, WA, USA [Internet]. Berkeley, CA: USENIX Association; 2016. Available: http://portal.acm.org/toc.cfm?id=1072430

10. Borger MA, Kaeding AF, Seeburger J, Melnitchouk S, Hoebartner M, Winkfein M, et al. Minimally invasive mitral valve repair in Barlow’s disease: Early and long-term results. The Journal of Thoracic and Cardiovascular Surgery. 2014;148: 1379–1385. doi:10.1016/j.jtcvs.2013.11.030

11. Coutinho GF, Antunes MJ. Mitral valve repair for degenerative mitral valve disease: Surgical approach, patient selection and long-term outcomes. Heart (British Cardiac Society). 2017;103: 1663–1669. doi:10.1136/heartjnl-2016-311031

12. Bianchi G, Margaryan R, Kallushi E, Cerillo AG, Farneti PA, Pucci A, et al. Outcomes of Video-assisted Minimally Invasive Cardiac Myxoma Resection. Heart, Lung and Circulation. 2017; doi:10.1016/j.hlc.2017.11.010

13. Expert Systems Research Science [Internet]. Available: http://science.sciencemag.org/content/220/4594/261.long

14. Holzhey DM, Seeburger J, Misfeld M, Borger MA, Mohr FW. Learning minimally invasive mitral valve surgery: A cumulative sum sequential probability analysis of 3895 operations from a single high-volume center. Circulation. 2013;128: 483–491. doi:10.1161/CIRCULATIONAHA.112.001402

